# Superficial stromal cell population in the mouse uterus require METTL14 for development and functional competence to support embryo implantation

**DOI:** 10.1101/2025.06.19.660542

**Authors:** Zhan-Hong Zheng, Run-Fu Jiang, Yu-Qi Hong, Qing-Yan Zhang, Yun-Tao Deng, Jiu-Qi Zhao, Jia-Peng He, Yan-Wen Xu, Shi-Hua Yang, Ji-Long Liu

**Affiliations:** College of Veterinary Medicine, South China Agricultural University, Guangzhou 510642, China; Jiangxi Provincial Key Laboratory for Animal Health, Institute of Animal Population Health, College of Animal Science and Technology, Jiangxi Agricultural University, Nanchang 330045, China; Department of Obstetrics and Gynecology, Reproductive Medical Center, The First Affiliated Hospital of Sun Yat-sen University, Guangzhou 510080, China; Guangdong Provincial Key Laboratory of Reproductive Medicine, the First Affiliated Hospital of Sun Yat-sen University, Guangzhou 510080, China

**Keywords:** METTL14, m^6^A, embryo implantation, estrogen receptor

## Abstract

Uterine stromal cells are critical for pregnancy, coordinating embryo implantation, placental development, and maintenance of gestation. Single-cell RNA-seq analyses have identified two distinct stromal cell populations in the mouse uterus: superficial and deep stromal cells. However, their functional specialization remains unresolved. Here, we show that mice lacking methyltransferase-like 14 (METTL14), a core enzyme of the m^6^A methyltransferase complex, exhibit a selective depletion of superficial stromal cells while preserving deep stromal cells. This genetic model allowed us to investigate the physiological role of superficial stromal cells in uterine function. Paralleled by the aberrant development and loss of superficial stromal cells, *Mettl14* deletion impaired embryo implantation and decidualization. Transcriptomic profiling revealed that *Mettl14* deficiency disrupts hormonal signaling by upregulating estrogen (E_2_)-responsive genes and downregulating progesterone (P_4_)-responsive genes, resulting in aberrant development and loss of superficial stromal cells. Mechanistically, *Mettl14* ablation reduces m^6^A methylation in the 3′-UTR of *Esr1* mRNA, thereby impairing m^6^A-dependent mRNA degradation and elevating ESR1 expression, which amplifies E_2_ signaling. This study underscores the essential role of METTL14 in maintaining the superficial stromal cell population required for embryo implantation by regulating *Esr1* mRNA stability through m^6^A modification.

## Introduction

The endometrium consists of two major cell types: epithelial cells and stromal cells. In the mouse embryo implantation model [1–3], the luminal epithelial cells proliferate and differentiate to establish receptivity, while stromal cells also contribute via paracrine signaling [4]. Glandular epithelial cells secrete leukemia inhibitory factor (LIF) to facilitate embryo attachment [5, 6]. Post-attachment, the epithelium induces stromal cell proliferation and differentiation (decidualization), through paracrine mechanisms [7, 8]. Following embryo invasion, decidualized stromal cells support placentation and pregnancy maintenance [9]. Glandular epithelium regulates post-implantation pregnancy by modulating stromal cell decidualization and placental development [10]. A deeper understanding of epithelial and stromal cell dynamics may inform strategies for addressing infertility and pregnancy-related disorders.

In recent years, single-cell RNA sequencing (scRNA-seq) has emerged as a powerful tool for dissecting cellular heterogeneity in complex tissues [11]. Using scRNA-seq, we identified two distinct stromal cell populations in the peri-implantation uterus: superficial stromal cells (SSCs), adjacent to the luminal epithelium, and deep stromal cells (DSCs), located deeper within the stroma [12–15]. SSCs proliferate on gestational day (GD) 4 [12] and continue proliferating post-attachment on GD5 [13], while both SSCs and DSCs proliferate following embryo invasion on GD6 [14]. Trajectory analysis revealed that SSCs form the primary decidual zone (PDZ), whereas DSCs adjacent to the PDZ contribute to the secondary decidual zone (SDZ) on GD8 [15]. A recent study demonstrated that SSCs and DSCs are present at birth [16], and analysis of the Mouse Cell Atlas (MCA) dataset confirmed their persistence in the adult uterus [12], suggesting their stability as distinct uterine layers. However, the functional roles of SSCs and DSCs remain poorly understood.

Here, we demonstrate that mice lacking METTL14, a core component of the m^6^A methyltransferase complex [17–19], exhibit selective depletion of SSCs while preserving DSCs, along with impaired embryo implantation and decidualization. Mechanistically, METTL14 deficiency disrupts hormonal signaling by upregulating estrogen (E_2_)-responsive genes and downregulating progesterone (P_4_)-responsive genes, leading to aberrant SSC development. Further investigation revealed that METTL14 ablation reduces m^6^A methylation in the 3′-UTR of *Esr1* mRNA, impairing m^6^A-dependent mRNA degradation and elevating ESR1 expression, thereby amplifying E_2_ signaling. Collectively, our findings establish METTL14 as critical for SSC development and function to support embryo implantation.

## Results

### *Pgr*-Cre-mediated deletion of *Mettl14* leads to pre-implantation embryo loss, defective uterine receptivity and compromised decidualization

METTL14 is highly expressed in both uterine epithelial and stromal cells (**Fig. S1**). Given that *Mettl14*-null mice are embryonically lethal [20], we generated conditional *Mettl14*- deleted mice (*Mettl14*^d/d^) by crossing *Mettl14*-floxed mice (*Mettl14*^f/f^) with *Pgr*-Cre driver mice (*Pgr*^Cre/+^) (**Fig. 1A**). PCR confirmed Cre-mediated excision of *Mettl14* exon 3 (flanked by loxP sites) in *Mettl14*^d/d^ uteri (**Fig. 1B**). RNA-seq revealed exon 3 deletion (88 bp) induced alternative splicing between exons 2 and 4, causing a frameshift (**Fig. 1C**). Exon 3-specific quantitative RT-PCR validated mRNA-level excision (**Fig. 1D**), while western blot and immunohistochemistry confirmed METTL14 protein loss (**Fig. 1E-F**). Dot blot analysis showed reduced global m^6^A RNA modification in *Mettl14*^d/d^ uteri compared to *Mettl14*^f/f^ (**Fig. 1G**), confirming efficient uterine METTL14 deletion. A 6-month fertility trial revealed infertility in *Mettl14*^d/d^ females but normal fertility in *Mettl14*^f/f^ females (**Fig. 1H**), demonstrating METTL14’s critical role in pregnancy.

**Figure 1.**
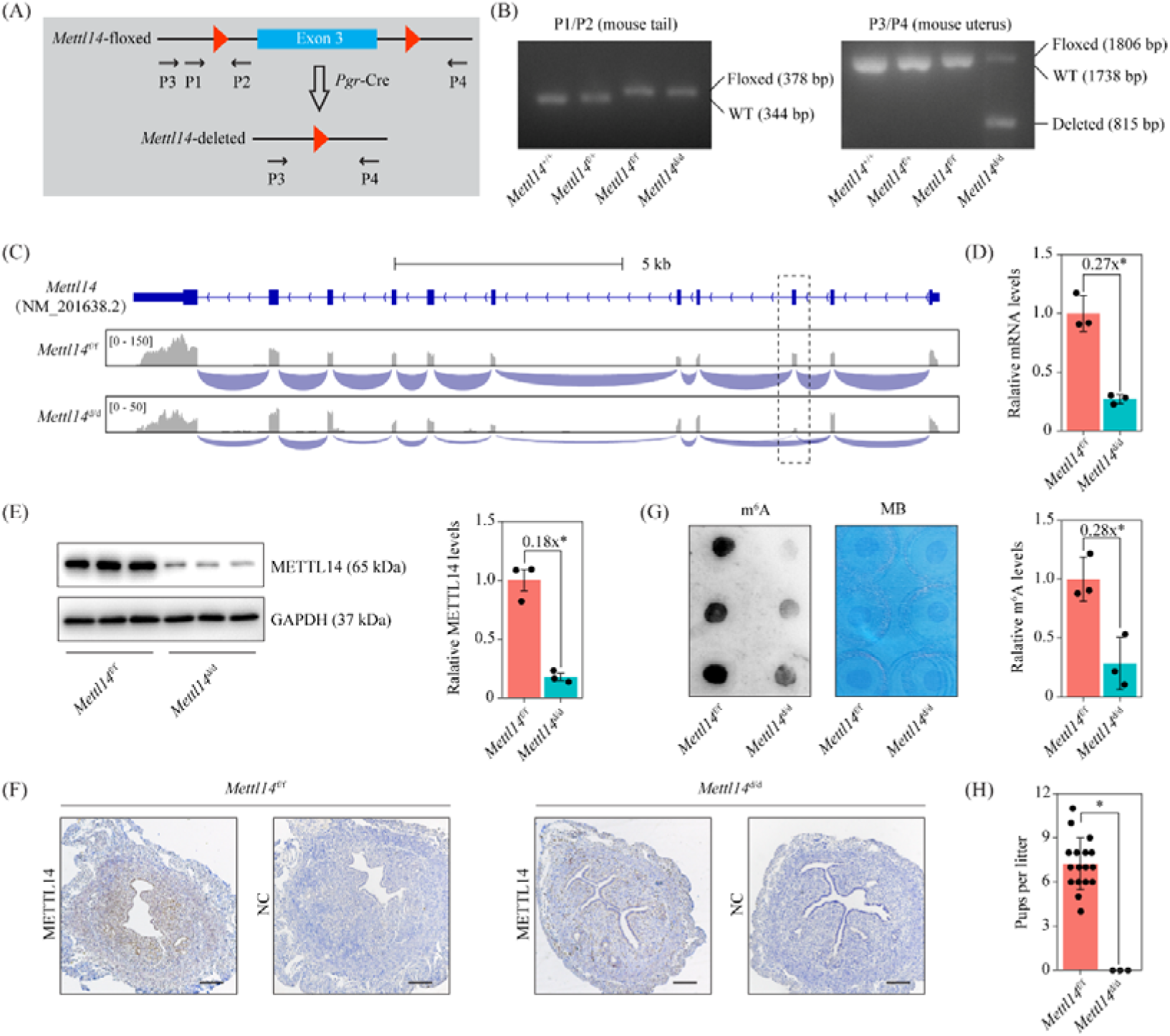
*Pgr*-Cre-mediated conditional deletion of *Mettl14* leads to female infertility in mice. (A) The diagram showing the strategy of conditional *Mettl14* deletion. (B) Genotyping analysis of *Mettl14*-floxed (*Mettl14*^f/f^) and *Mettl14*-deleted (*Mettl14*^f/f^*Pgr*^Cre/+^; *Mettl14*^d/d^) mice. (C) Integrative Genomics Viewer showing the deletion of exon 3 of *Mettl14* in the uterus of *Mettl14*^d/d^ mice based on RNA-seq analysis. (D) Quantitative RT-PCR analysis of *Mettl14* mRNA levels at exon 3 in the uterus of *Mettl14*^f/f^ and *Mettl14*^d/d^ mice on GD4. Data are presented as mean ± SD. *, P < 0.05. (E) Western blot analysis of METTL14 protein levels in the uterus of *Mettl14*^f/f^ and *Mettl14*^d/d^ mice on GD4. Data are presented as mean ± SD. *, P < 0.05. (F) Immunohistochemistry staining of METTL14 protein in the uterus on GD4. NC, no primary antibody control. Bar = 100 μm. (G) Dot blot of m^6^A levels in the uterus of of *Mettl14*^f/f^ and *Mettl14*^d/d^ mice on GD4. MB, methylene blue. Data are presented as mean ± SD. *, P < 0.05. (H) Bar plot showing litter sizes for *Mettl14*^f/f^ mice and *Mettl14*^d/d^ mice during a 6-month fertility test. Data are presented as mean ± SD. n=3 mice. *, P < 0.05.

To elucidate the cause of infertility in *Mettl14*^d/d^ mice, we assessed their pregnancy status at various timepoints. *Mettl14*^d/d^ mice showed no implantation sites on GD8 (**Fig. 2A**), a finding traceable back to GD5 (**Fig. 2B**). We detected a complete loss of pre-implantation embryos in the reproductive tract on GD4 (**Fig. 2C**), with an earlier loss evident on GD3 (**Fig. 2D**). However, embryos were recovered from the oviduct of *Mettl14*^d/d^ mice on GD2 and could be cultured to blastocysts in vitro (**Fig. 2E**). Immunohistochemistry revealed unaffected METTL14 expression in *Mettl14*^d/d^ mouse ovaries on GD4, with normal expression of the steroid biosynthetic enzyme CYP11A1 in the corpus luteum (**Fig. 2F**). Serum levels of P_4_ and E_2_ were comparable between *Mettl14*^d/d^ and *Mettl14*^f/f^ mice on GD4 (**Fig. 2G**). Examining the oviduct, we found intact METTL14 in the ampulla but efficient deletion in the isthmus (**Fig. 2H**). Taken together, these findings suggest that the deletion of *Mettl14* in the isthmus creates a hostile oviduct environment, which is responsible for the pre-implantation embryo loss in *Mettl14*^d/d^ mice.

**Figure 2.**
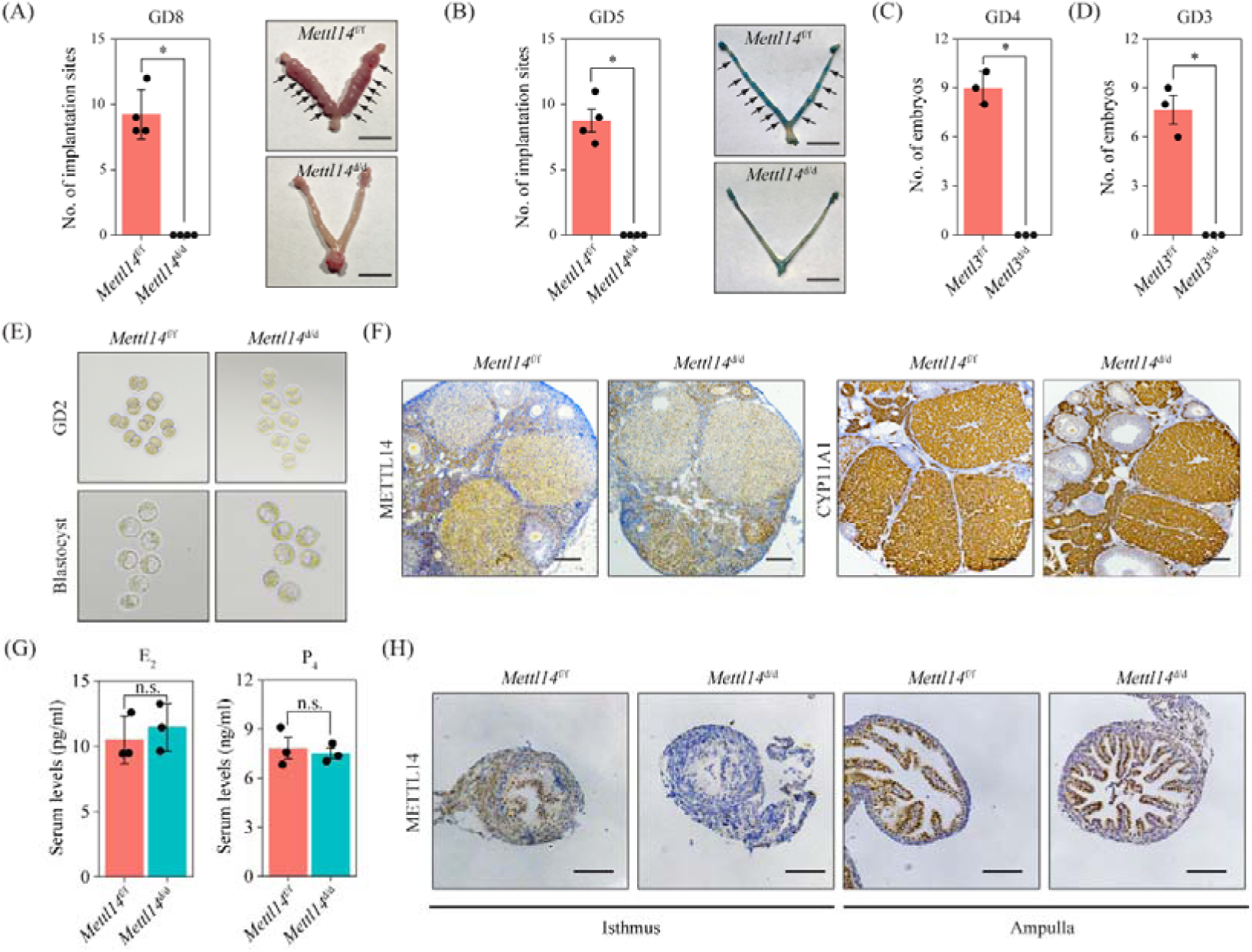
*Mettl14* deletion results in implantation failure due to embryo loss in the oviduct. (A-B) Bar plot showing the number of embryo implantation sites in *Mettl14*^f/f^ and *Mettl14*^d/d^ uteri on GD8 (A) and GD5 (B). Implantation sites are marked by arrowheads. Scale bar = 1 cm. Data are presented as mean ± SD. *, P < 0.05. (C-D) Bar plot showing the number of embryos collected from the uterus on GD4 (C) and the oviduct on GD3 (D). Data are presented as mean ± SD. *, P < 0.05. (E) Images of embryos flushed from the oviduct on GD2 (upper panel) and images of in-vitro cultured blastocysts (lower panel) from *Mettl14*^f/f^ and *Mettl14*^d/d^ mice. (F) Immunohistochemical staining of METTL14 protein and CYP11A1 protein in the ovary of *Mettl14*^f/f^ and *Mettl14*^d/d^ mice on GD4. Scale bar = 100 μm. (G) The concentration of circulating E_2_ and P_4_ in *Mettl14*^f/f^ and *Mettl14*^d/d^ mice on GD4. Data are presented as mean ± SD. n.s., not significant. (H) Immunohistochemical staining of METTL14 protein in the oviduct of *Mettl14*^f/f^ and *Mettl14*^d/d^ mice on GD3. Scale bar = 100 μm.

To assess uterine receptivity, we performed embryo transfer experiments. Blastocysts from *Mettl14*^d/d^ donors successfully implanted in *Mettl14*^f/f^ recipients, while those from *Mettl14*^f/f^ donors failed in *Mettl14*^d/d^ recipients (**Fig. 3A**). Immunohistochemical analysis of MKI67 indicated persistent LE proliferation in *Mettl14*^d/d^ mice on GD4 (**Fig. 3B**). MUC1, an indicator of uterine receptivity, was abundant on the LE of *Mettl14*^d/d^ mice but absent on that of *Mettl14*^f/f^ mice on GD4 (**Fig. 3C**). CDH1, a marker of cell junction and polarity, was more intensely expressed in the LE of *Mettl14*^d/d^ mice (**Fig. 3D**). Scanning and transmission electron microscopy revealed a lack of pinopodes and dense microvilli on the apical LE of *Mettl14*^d/d^ uteri, in contrast to *Mettl14*^f/f^ uteri (**Fig. 3D-E**). These results demonstrate defective uterine receptivity in *Mettl14*^d/d^ mice.

**Figure 3.**
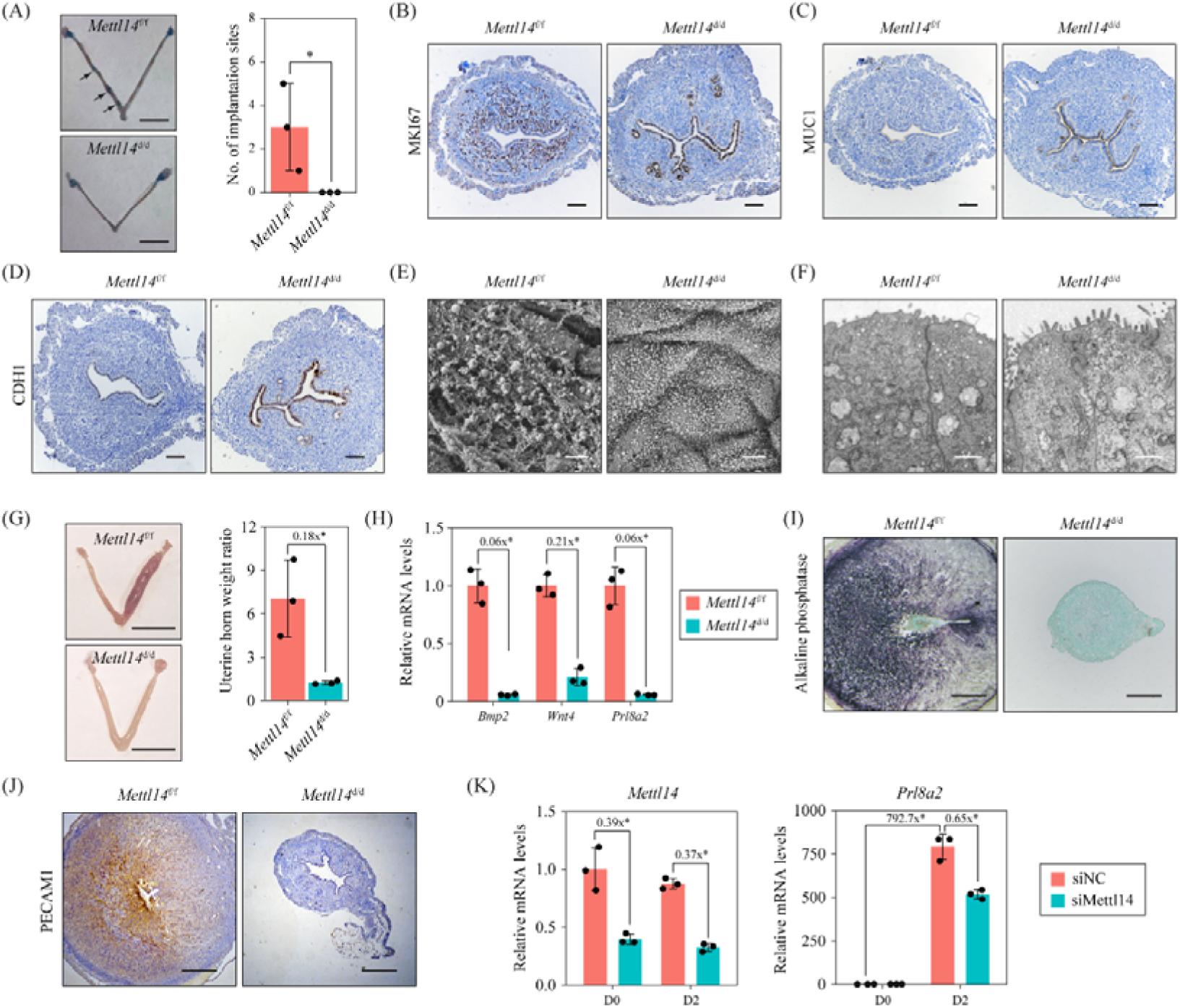
*Mettl14* deletion causes impaired uterine receptivity and defective decidual response. (A) Embryo transfer experiment between *Mettl14^f/f^* and *Mettl14^d/d^*mice. Six cultured blastocysts recovered from the oviduct of *Mettl14^f/f^* mice on GD2 were transferred to one uterine horn of pseudo-pregnant *Mettl14^d/d^*mice, and vice versa. Embryo implantation sites were examined on GD6. Data are presented as mean ± SD. *, P < 0.05. (B-D) Immunohistochemistry staining of MKI67 (B), MUC1 (C) and CDH1 (D) in *Mettl14*^f/f^ and *Mettl14*^d/d^ uteri on GD4. Scale bar = 100 μm. (E) Scanning electron microscopy analysis of the luminal surface of *Mettl14*^f/f^ and *Mettl14*^d/d^ uteri on GD4. Scale bar = 2 μm. (F) Transmission electron microscopic analysis of the luminal epithelial cells of *Mettl14*^f/f^ and *Mettl14*^d/d^ uteri on GD4. Scale bar = 1 μm. (G) Artificial decidualization in *Mettl14*^f/f^ and *Mettl14*^d/d^ mice. The weight ratio between the stimulated horn and the un-stimulated horn was calculated. Data are presented as mean ± SD. *, P < 0.05. (H) Quantitative RT-PCR analysis of decidualization marker genes in the stimulated uterine horn of *Mettl14*^f/f^ and *Mettl14*^d/d^ mice. Data are presented as mean ± SD. *, P < 0.05. (I) Alkaline phosphatase activity staining of the stimulated uterine horn from *Mettl14*^f/f^ and *Mettl14*^d/d^ mice. Bar = 500 μm. (E) Immunohistochemistry staining of PECAM1 in the stimulated uterine horn from *Mettl14*^f/f^ and *Mettl14*^d/d^ mice. Bar = 500 μm. (K) Quantitative RT-PCR analysis of *Prl8a2* expression in in-vitro decidualized primary mouse endometrial stromal cells after *Mettl14* knockdown. Data are presented as means ± SD. *, P < 0.05.

To investigate the impact of *Mettl14* deletion on decidualization, we employed an artificial decidualization model. *Mettl14*^f/f^ mice exhibited a robust decidual response, absent in *Mettl14*^d/d^ mice (**Fig. 3G**). Quantitative RT-PCR analysis of decidualization marker genes (*Bmp2*, *Wnt4*, *Prl8a2*) confirmed compromised decidualization in *Mettl14*^d/d^ mice (**Fig. 3H**). Alkaline phosphatase activity, an indicator of stromal cell differentiation, was evident in *Mettl14*^f/f^ but not *Mettl14*^d/d^ uteri post-decidualization (**Fig. 3I**). Immunostaining for PECAM1, a marker of endothelial cells, revealed reduced angiogenesis in *Mettl14*^d/d^ uteri after artificial decidualization (**Fig. 3J**). In vitro decidualization of primary mouse endometrial stromal cell (mESCs) from wild-type mice showed impaired *Prl8a2* induction upon METTL14 knockdown using siRNA (**Fig. 3K**). In summary, *Mettl14* deletion leads to compromised decidualization.

### *Mettl14* deletion causes a loss of superficial stromal cells (SSCs)

To gain an understanding of the cell type-specific alternations in *Mettl14*^d/d^ uteri, we conducted single-cell RNA-seq analysis on GD4. After quality control, a total of 10098 cells (5332 cells for *Mettl14*^f/f^ uteri and 4766 cells for *Mettl14*^d/d^ uteri) were obtained (**Fig. 4A**). Unsupervised clustering analysis revealed 12 distinct cell types defined by the expression of known marker genes as we described previously vv: luminal epithelial cells (LE, TACSTD2^+^), glandular epithelial cells (GE, FOXA2^+^), superficial stromal cells (SSC, PGR^+^VIM^+^ACTA2^-^HAND2^high^), deep stromal cells (DSC, PGR^+^VIM^+^ACTA2^-^HAND2^low^); smooth muscle cells (SMC, ACTA2^+^RGS5^-^); pericytes (PC, ACTA2^+^RGS5^+^), vascular endothelial cells (VEC, PECAM1^+^VWF^+^), lymphatic endothelial cells (LEC, PECAM1^+^PROX1^+^), natural killer cells (PTPRC^+^NKG7^+^), T cells (PTPRC^+^CD3E^+^), macrophages (M, PTPRC^+^ADGRE1^+^), and dendritic cells (DC, PTPRC^+^CD209A^+^) (**Fig. 4B**).

**Figure 4.**
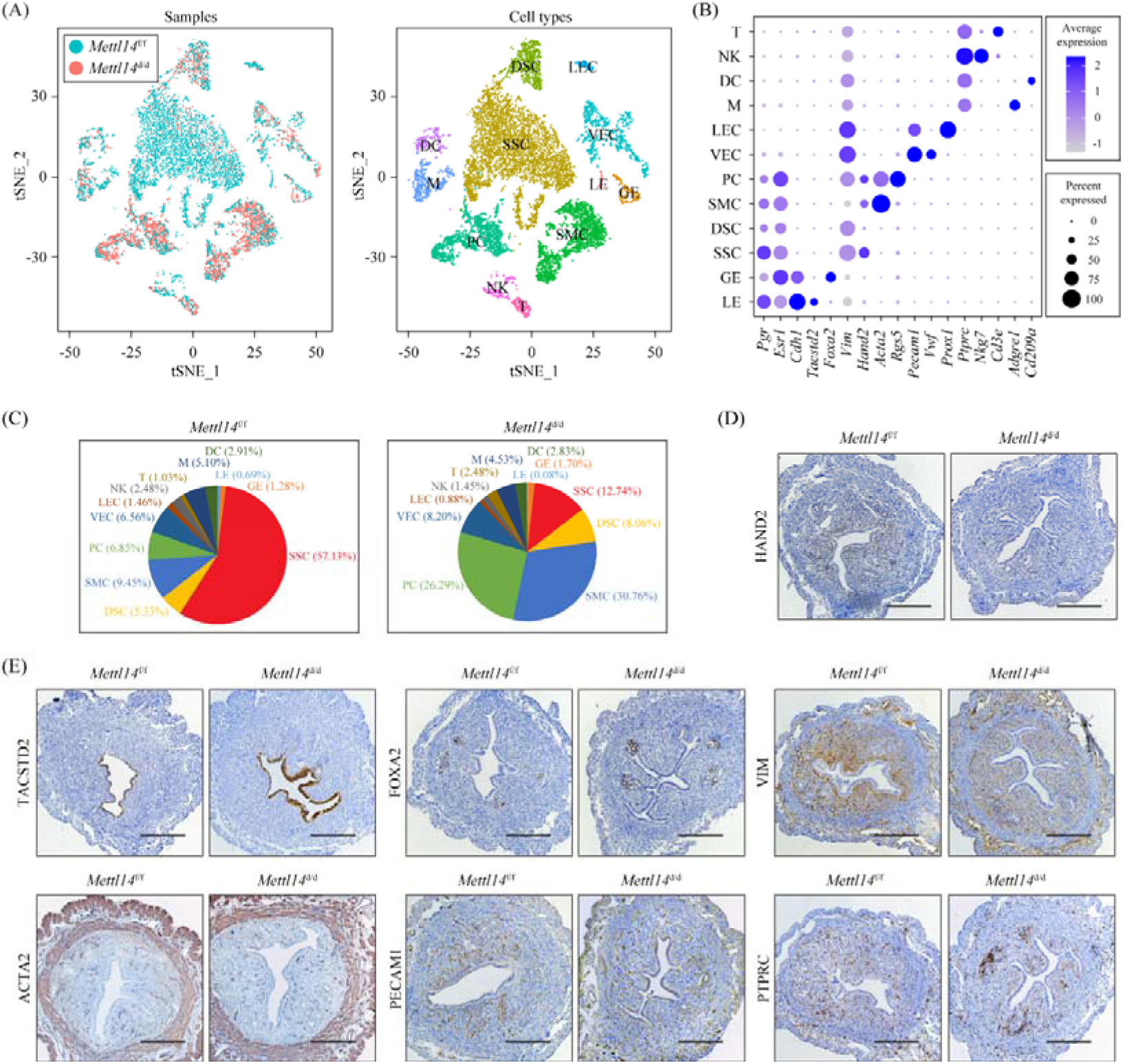
The number of superficial stromal cells is significantly decreased in the *Mettl14*-deleted uteri during the window of implantation. (A) ) TSNE visualization of single-cell RNA-seq data from *Mettl14*^f/f^ and *Mettl14*^d/d^ uteri. Single cells were grouped by samples (left panel) and cell clusters (right panel), respectively. LE, luminal epithelial cells; GE, glandular epithelial cells; SSC, superficial stromal cells; DSC, deep stromal cells; SMC, smooth muscle cells; PC, pericytes; VEC, vascular endothelial cells; LEC, lymphatic endothelial cells; NK, natural killer cells; T, T cells; M, macrophages; DC, dendritic cells. (B) Dot plot showing the expression pattern of canonical marker genes for different cell types. (C) Pie plot showing cell population change of 12 major cell types in *Mettl14*^f/f^ and *Mettl14*^d/d^ uteri. (D) Immunohistochemistry staining of HAND2 in *Mettl14*^f/f^ and *Mettl14*^d/d^ uteri on GD4. Bar = 100 μm. (E) Immunohistochemistry staining of LE marker TACSTD2, GE marker FOXA2, mesenchymal cell marker VIM, SMC/PC marker ACTA2, and immune cell marker PTPRC in *Mettl14*^f/f^ and *Mettl14*^d/d^ uteri on GD4. Bar = 100 μm.

We then examined the abundance of each cell type and found that the proportion of SSCs was significantly decreased, while the proportions of SMCs and PCs were significantly increased (**Fig. 4C**). To further validate these findings, we used immunostaining techniques targeting specific markers for each cell type, which revealed a notable reduction in the absolute number of SSCs (**Fig. 4D**), whereas the absolute counts of SMCs and PCs seemed unaltered (**Fig. 4E**). Thus, the observed increase in the proportions of SMCs and PCs in our single-cell RNA-seq data can be attributed to the decrease in SSC cell numbers, resulting in a reduced total cell count and, consequently, an elevated percentage of other cell types. We then performed differential gene expression analysis in each cell type with a fold change cut[off of 1.5 and a P value cut[off of 0.05 (**Table S1**). Based on the number of differentially expressed genes, we found that the SSC is the most affected cell type in *Mettl14*^d/d^ uteri (**Fig. S2A**). In the SSC, we found that a total of 1404 genes were differentially expressed, of which 774 genes were downregulated and 630 genes were upregulated in *Mettl14*^d/d^ uteri in comparison to *Mettl14*^f/f^ uteri (**Fig. S2B**). GO analysis revealed that the downregulated genes were notably associated with decidualization (**Fig. S2C**). Analysis of cell-cell communication revealed a loss of interaction between LE and other cells in *Mettl14*^d/d^ uteri compared to *Mettl14*^f/f^ uteri (**Fig. S3A**). Notably, FGF1/2/7/10/22 were expressed in SSC, while their corresponding receptor FGFR2 was expressed in LE/GE (**Fig. S3B**), emphasizing the importance of the FGF signaling pathway. This pathway integrates with various other signaling cascades, such as Calcium, Rap1, Ras, PI3K-Akt, and MAPK signaling pathways (**Fig. S3C**). Collectively, our findings suggest that the reduced population of SSCs in *Mettl14*^d/d^ uteri represents a critical defect.

### *Mettl14* deletion causes elevated E_2_ signaling due to the upregulation of ESR1

To elucidate the downstream effector of METTL14, we conducted RNA-seq analysis to compare the gene expression profiles of *Mettl14*^d/d^ and *Mettl14*^f/f^ uteri on GD4 (**Fig. 5A**). The analysis identified 1472 differentially expressed genes (fold change > 2 and adjusted P < 0.05), comprising 835 upregulated and 637 downregulated genes in *Mettl14*^d/d^ uteri compared to *Mettl14*^f/f^ uteri (**Table S2**). Gene ontology (GO) analysis revealed that the upregulated genes were notably associated with inflammatory responses (**Fig. S4**). We noticed that P_4_ target genes, including heart and neural crest derivatives expressed 2 (*Hand2*), homeobox A10 (*Hoxa10*), Indian hedgehog (*Ihh*) and amphiregulin (*Areg*) were significantly down-regulated, while E_2_ target genes, including mucin 1 (*Muc1*), mucin 4 (*Muc4*), complement C3 (*C3*) and lactotransferrin (*Ltf*) were significantly upregulated in *Mettl14*^d/d^ uteri compared to *Mettl14*^f/f^ uteri. These alterations were further validated through quantitative RT-PCR using an independent set of uterine samples (**Fig. 5B**). Additionally, quantitative RT-PCR demonstrated that *Esr1* but not *Pgr* was significantly upregulated in *Mettl14*^d/d^ uteri compared to *Mettl14*^f/f^ uteri (**Fig. S5A-B**). Western blot (**Fig. 5C**) and immunohistochemical staining (**Fig. 5D**) further corroborated this finding, revealing increased levels of ESR1 and phosphorylated ESR1 (p-ESR1), while PGR expression remained unaffected in *Mettl14*^d/d^ uteri. Collectively, these results point to altered E_2_ and P_4_ signaling resulting from an increased expression of ESR1 in *Mettl14*^d/d^ uteri.

**Figure 5.**
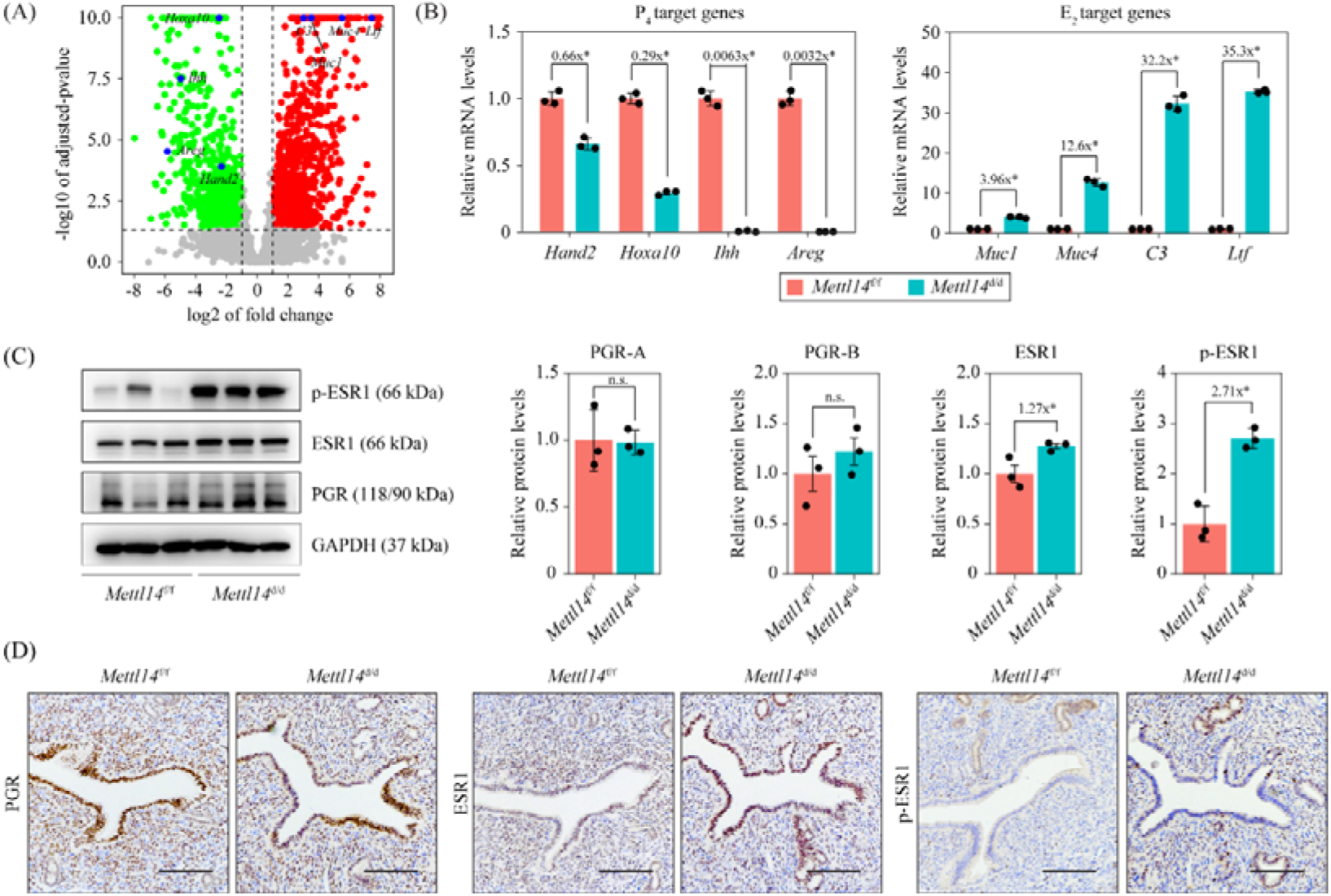
ESR1 is significantly upregulated in *Mettl14*-deleted uteri during the window of implantation. (A) Volcano plot showing uterine gene expression profiles of *Mettl14*^f/f^ and *Mettl14*^d/d^ uteri on GD4. Differentially expressed genes (fold change > 2 and adjusted P < 0.05) are denoted in red or green. (B) Validation of P_4_ target genes and E_2_ target genes by using quantitative RT-PCR. Data are presented as mean ± SD. *, P < 0.05. (C) Western blot of PGR, ESR1 and p-ESR1 in *Mettl14*^f/f^ and *Mettl14*^d/d^ uteri on GD4. Data are presented as mean ± SD. *, P < 0.05. (D) Immunohistochemistry staining of PGR, ESR1 and p-ESR1 in *Mettl14*^f/f^ and *Mettl14*^d/d^ uteri on GD4. Bar = 100 μm.

### *Esr1* mRNA is a direct target of m^6^A modification

To further explore the global m^6^A modification on mRNAs in the uteri of wild-type mice on GD4, we utilized methylated RNA immunoprecipitation sequencing (MeRIP-seq). In total, 24127 m^6^A peaks, corresponding to 14412 genes, were identified (**Table S3**). Notably, a significant portion of these peaks (94.97%) mapped to protein-coding genes (**Fig. 6A**). Among the protein-coding genes with m^6^A peaks, the majority (76.03%) resided in the coding sequence (CDS), followed by the intronic region (11.51%), the 3′-untranslated region (UTR) (8.34%), and finally, the 5′-UTR (4.12%) (**Fig. 6B**). Metagene analysis revealed a pronounced enrichment of m^6^A peaks around start and stop codons in mature mRNAs (**Fig. 6C**). An unbiased motif search revealed that the RRACH motif (R = A/G, H = A/C/U) was significantly enriched in m^6^A peaks (**Fig. 6D**), consistent with previous findings [21]. Gene ontology (GO) enrichment analysis indicated that m^6^A modification was most strongly associated with transcription (**Fig. 6E**). To examine the effect of m^6^A modification on gene expression, we re-analyzed RNA-seq data from *Mettl14*^d/d^ and *Mettl14*^f/f^ uteri on GD4. Differentially expressed genes were then categorized as m^6^A-positive or m^6^A-negative based on the MeRIP-seq data. Interestingly, we observed a small but statistically significant increase in relative mRNA abundance for m^6^A-positive genes following *Mettl14* deletion (**Fig. 6F**). Furthermore, m^6^A modification was slightly but significantly more prevalent among upregulated genes (55.63%) compared to downregulated genes (50.47%) in *Mettl14*^d/d^ uteri relative to *Mettl14*^f/f^ uteri (**Fig. 6G**). These findings align with previous studies suggesting that m^6^A modification in the uterus primarily functions to promote mRNA degradation.

**Figure 6.**
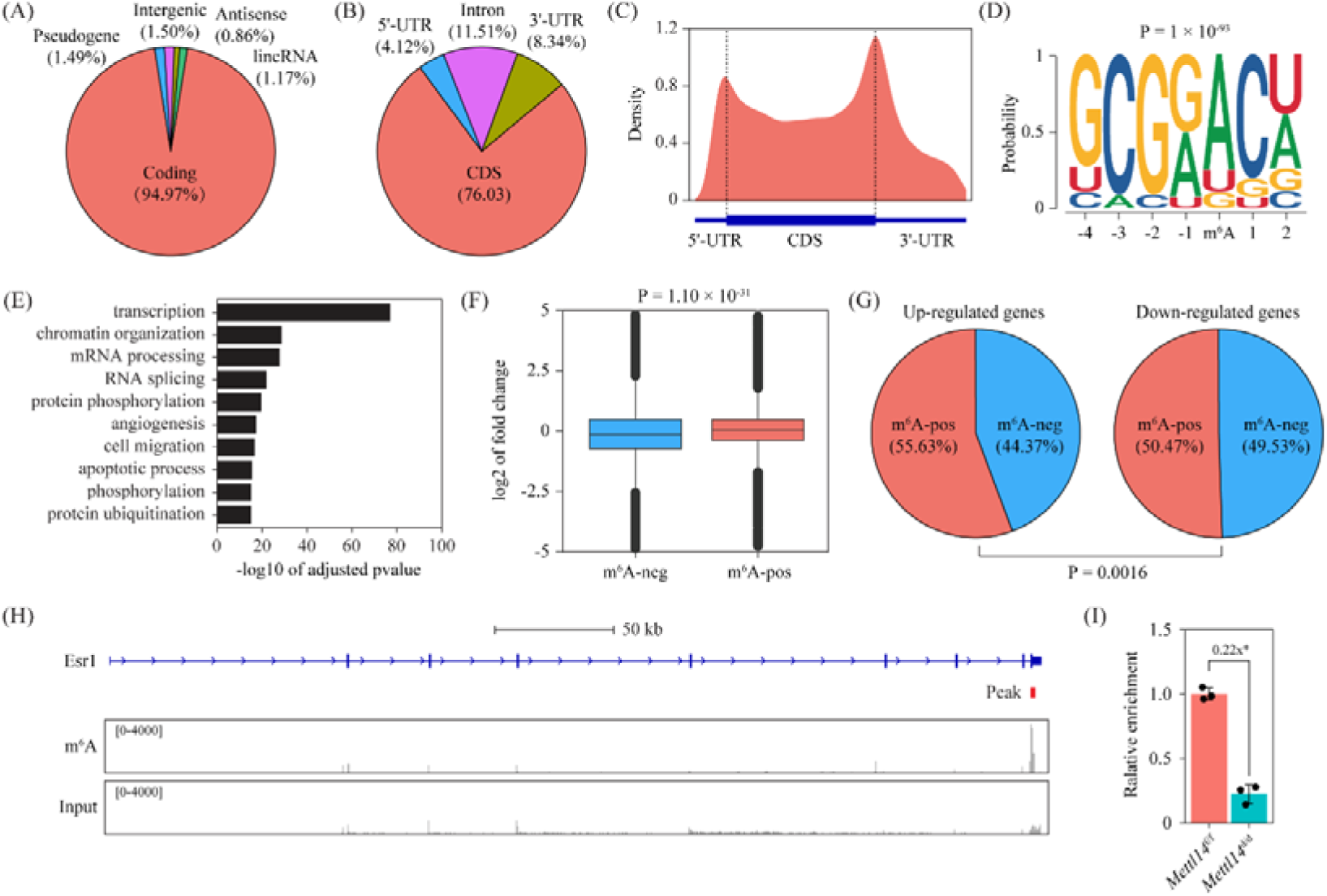
MeRIP-seq analysis reveals that *Esr1* mRNA is a direct target of METTL14- mediated m^6^A modification. (A-E) Profiling of global m^6^A modification in mouse uterus on GD4 using MeRIP-seq. (A) Pie chart presenting fractions of m^6^A peaks in different genomic segments. (B) Pie chart presenting fractions of m^6^A peaks within coding genes. (C) The metagene distribution of m^6^A peaks in coding genes. (D) Sequence logo of the enriched motif in m^6^A peaks. (E) Bar plot showing the top 10 most enriched gene ontology terms among m^6^A-positive genes ranked by adjusted P-value. (F-G) Comparing the expression trend of m6A-positive genes and m6A-negtive genes in the uterus on GD4. (F) Box plots comparing log2 of fold change values in m^6^A-positive genes and m^6^A-negtive genes. The P-value is calualted by Wilcoxon rank-sum test. (G) Pie plots showing the percentage of m^6^A positive genes in up-regulated genes and down-regulated genes in *Mettl14*^d/d^ uteri compared to *Mettl14*^f/f^ uteri. (H) Integrative Genomics Viewer displaying MeRIP-seq coverage data at the *Esr1* gene locus. Peak calling was performed by using the MACS3 software. The most signifanct peak based on adjusted P-value is shown. (I) Validation of *Esr1* m^6^A modification by quantitative MeRIP-PCR. Data are presented as means ± SD. *, P < 0.05.

By MeRIP-seq data mining, we identified an m^6^A peak in the 3′-UTR of *Esr1* mRNA (**Fig. 6H**). Quantitative MeRIP-PCR analysis revealed a significantly decreased enrichment of *Esr1* mRNA in the uterus of *Mettl14*^d/d^ mice compared to *Mettl14*^f/f^ mice on GD4 (**Fig. 6I**). These results indicate that *Esr1* mRNA is a direct target for METTL14-mediated m^6^A modification in mouse uterus.

### METTL14-dependent m^6^A modification in the 3**′** UTR of *Esr1* mRNA promotes degradation

The m^6^A modification exerts a contextual regulation on gene expression, encompassing effects on mRNA splicing, translocation, degradation, stabilization, and translation. Given the observed upregulation of *Esr1* mRNA following *Mettl14* deletion in mouse uteri, we postulated that m^6^A modification of *Esr1* mRNA functions to enhance its degradation. To validate this hypothesis, we isolated primary mouse endometrial stromal cells (mESCs) from wild-type mice on GD4. Knockdown of *Mettl14* via siRNA significantly upregulated *Esr1* mRNA expression in these cells (**Fig. 7A**). Subsequently, we examined the influence of METTL14 on the stability of *Esr1* mRNA in MESCs. Upon treatment with actinomycin-D to block transcription, we observed a significant increase in the stability of *Esr1* mRNA at 12 h following *Mettl14* knockdown (**Fig. 7B**). These findings substantiate our hypothesis that *Mettl14*-dependent m^6^A modification serves to promote the degradation of *Esr1* mRNA.

**Figure 7.**
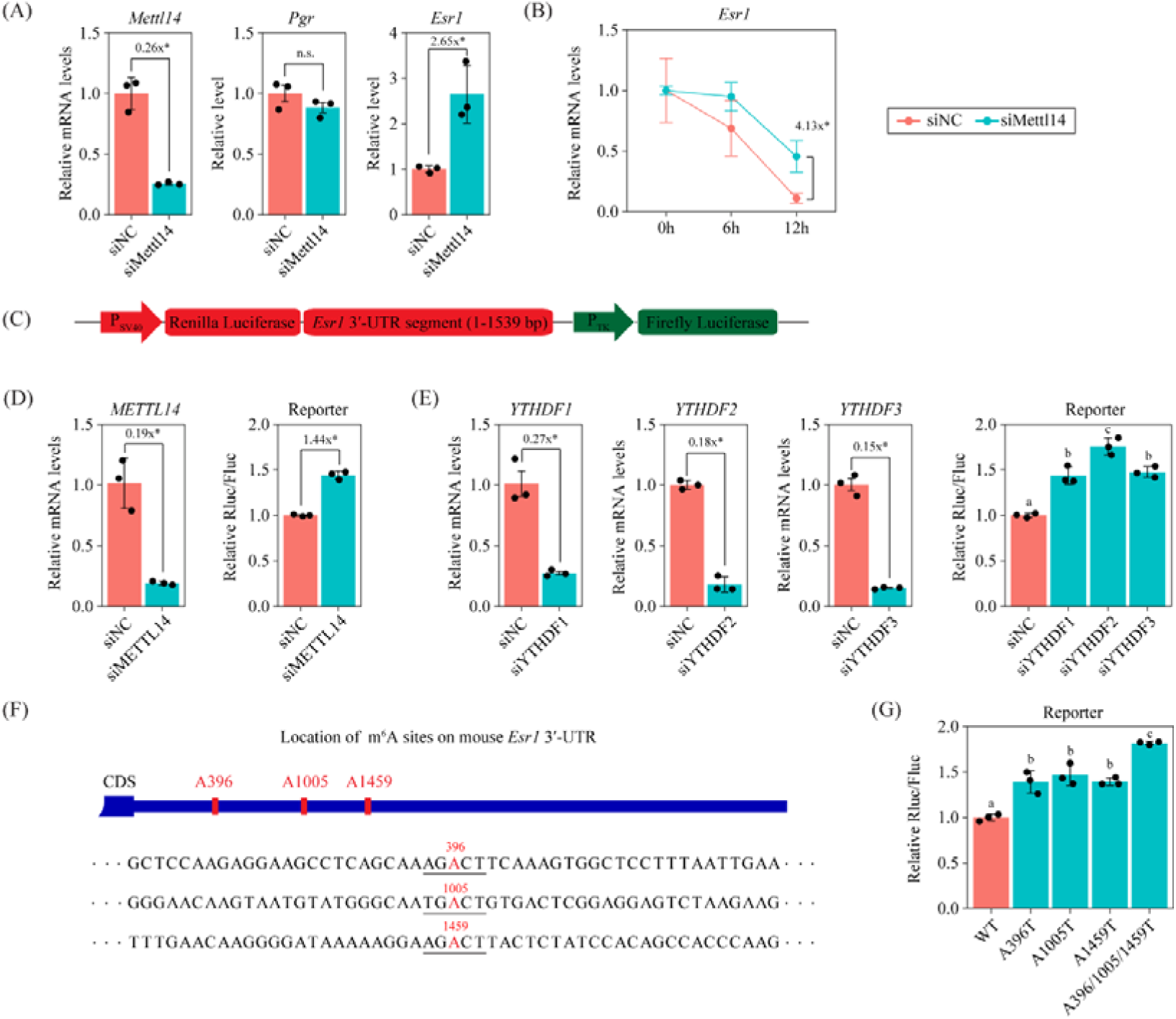
**The m^6^A modification in the 3**′**-UTR of *Esr1* mRNA promotes degradation.** (A) Quantitative RT-PCR analysis of *Pgr* and *Esr1* mRNA levels in primary mouse endometrial stromal cells (MESCs) after *Mettl14* knockdown for 48 h. Data are presented as means ± SD. *, P < 0.05. (B) Quantitative RT-PCR analysis of *Esr1* mRNA levels in MESCs after the treatment of actinomycin D at indicated times. Data are presented as means ± SD. *, P < 0.05. (C) Mouse *Esr1* 3′-UTR reporter for dual-luciferase assay. (D) Dual-luciferase assay in HEK293T cells after *Mettl14* knockdown. Data are presented as means ± SD. *, P < 0.05. (E) Dual-luciferase assay in HEK293T cells after *YTHDF1/2/3* knockdown. Data are presented as means ± SD. *, P < 0.05. (F) The location of m^6^A sites in the 3′-UTR of *Esr1* mRNA predicted by the SRAMP tool. Potential m^6^A sites are colored in red and the corresponding RRACH motifs are underlined. (G) Dual-luciferase assay in HEK293T cells using A-to-T point mutants. Data are presented as means ± SD. Different letters above the bar indicate statistically significant difference at P < 0.05.

To delve deeper into the role of m^6^A modification within the 3′-UTR of *Esr1* mRNA, we generated a luciferase reporter construct, incorporating a segment of the mouse *Esr1* 3′-UTR sequence into the 3′-UTR of *Renilla* luciferase (**Fig. 7C**). To ensure transfection consistency, we used firefly luciferase as a control. This luciferase reporter was subsequently co-transfected into HEK293 cells, along with *METTL14* siRNA or a negative control siRNA. Our results revealed a significant increase in relative luciferase activity upon *METTL14* knockdown (**Fig. 7D**). Previous studies have identified YTHDF2, YTHDF3, and, to a lesser extent, YTHDF1 as m^6^A readers that promote the degradation of m^6^A-modified mRNAs [17–19]. To determine which of these m^6^A readers mediates this regulatory process, we employed siRNA-mediated knockdown of these proteins. As illustrated in **Fig. 7E**, knockdown of either YTHDF1, YTHDF2, or YTHDF3 led to a significant enhancement in relative luciferase activity. These findings suggest that m^6^A modification in the 3′-UTR of *Esr1* mRNA enhances its degradation in a YTHDF1/2/3-dependent manner.

To precisely identify the location of m^6^A modification sites in *Esr1* mRNA, we employed the SRAMP prediction tool. This analysis pinpointed three potential m^6^A sites: A85, A310, and A317 (**Fig. 7F**). To functionally validate these predictions, we introduced A-to-T point mutations into the luciferase reporter plasmid harboring the wild-type *Esr1* 3′-UTR, thereby abrogating each of the identified m^6^A modification sites. Through luciferase assays, we observed that the A85T, A310T, and A317T mutations individually led to a significant enhancement in relative luciferase activity compared to the wild-type control (**Fig. 7G**). Notably, the triple mutation (A85T/A310T/A317T) exhibited an even more profound effect, suggesting that these three m^6^A modification sites likely function in a synergistic manner.

### METTL14 is required for human endometrial stromal cell (hESC) decidualization in vitro

In primary human endometrial stromal cells (hESCs), we observed an upregulation of *ESR1* mRNA expression upon *METTL14* knockdown using siRNA (**Fig. 8A**). To gain further insights, we conducted an RNA decay assay and found that the relative *ESR1* mRNA level increased following *METTL14* knockdown after Actinomycin-D treatment for 6-12 h (**Fig. 8B**). These results indicated that the METTL14-ESR1 axis is conserved between mice and humans. To study the function of METTL4 in human decidualization, we utilized a well-established in vitro model of decidualization, wherein hESCs undergo the decidualization process upon stimulation with 1 μM MPA and 0.5 mM 8-Br-cAMP. Our quantitative RT-PCR (**Fig. 8C**) and western blot (**Fig. 8D**) analyses revealed a significant downregulation in the expression of key decidualization marker genes, prolactin (PRL) and insulin-like growth factor binding protein 1 (IGFBP1), upon *METTL14* knockdown after 4 days of decidualization. These findings demonstrate that METTL14 is indispensable for the decidualization of hESCs in vitro.

**Figure 8.**
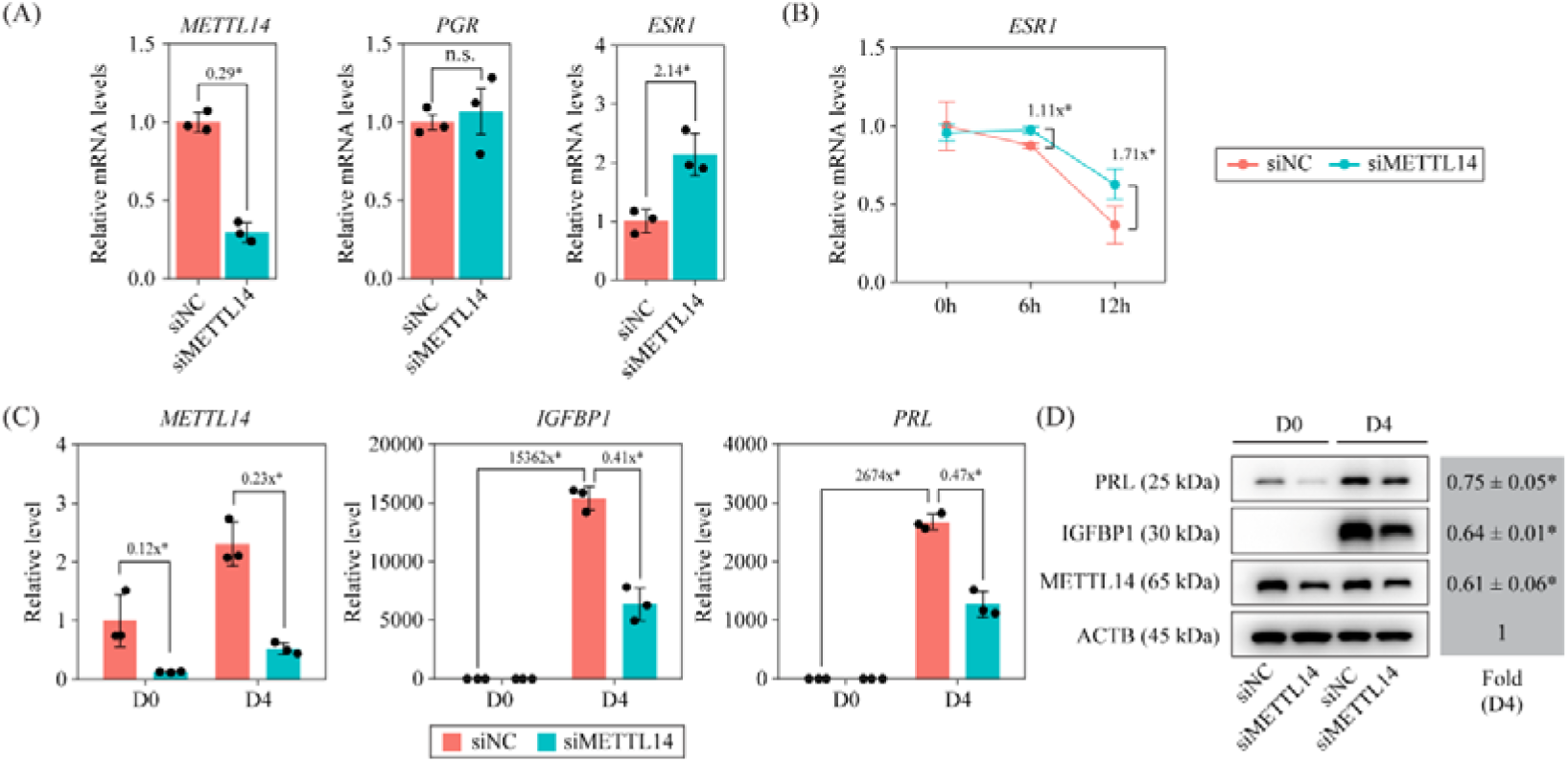
METTL14 is required for human endometrial stromal cell (hESC) decidualization in vitro. (A) Quantitative RT-PCR analysis of *PGR* and *ESR1* mRNA levels in primary hESCs after *METTL14* knockdown for 48 h. Data are presented as means ± SD. *, P < 0.05. (B) Quantitative RT-PCR analysis of *ESR1* mRNA levels in hESCs after the treatment of actinomycin D at indicated times. Data are presented as means ± SD. *, P < 0.05. (C) Quantitative RT-PCR analysis of *PRL* and *IGFBP1* mRNA levels in hESCs following in vitro decidualization. D0, day 0 of decidualization. D4, day 4 of decidualization. Data are presented as means ± SD. *, P < 0.05. (D) Western blot analysis of PRL and IGFBP1 protein levels in hESCs following in vitro decidualization. Data are presented as means ± SD. *, P < 0.05.

## Discussion

Previously, we characterized two distinct uterine stromal cell populations in mice: superficial stromal cells (SSCs) and deep stromal cells (DSCs) [12–15]. In this study, we demonstrate that uterine-specific knockout of METTL14 in mice leads to SSC loss. This model provides a valuable tool to elucidate the physiological roles of SSCs in the uterus and to uncover the molecular mechanisms underlying their normal development and function (**Fig. 9**).

**Figure 9.**
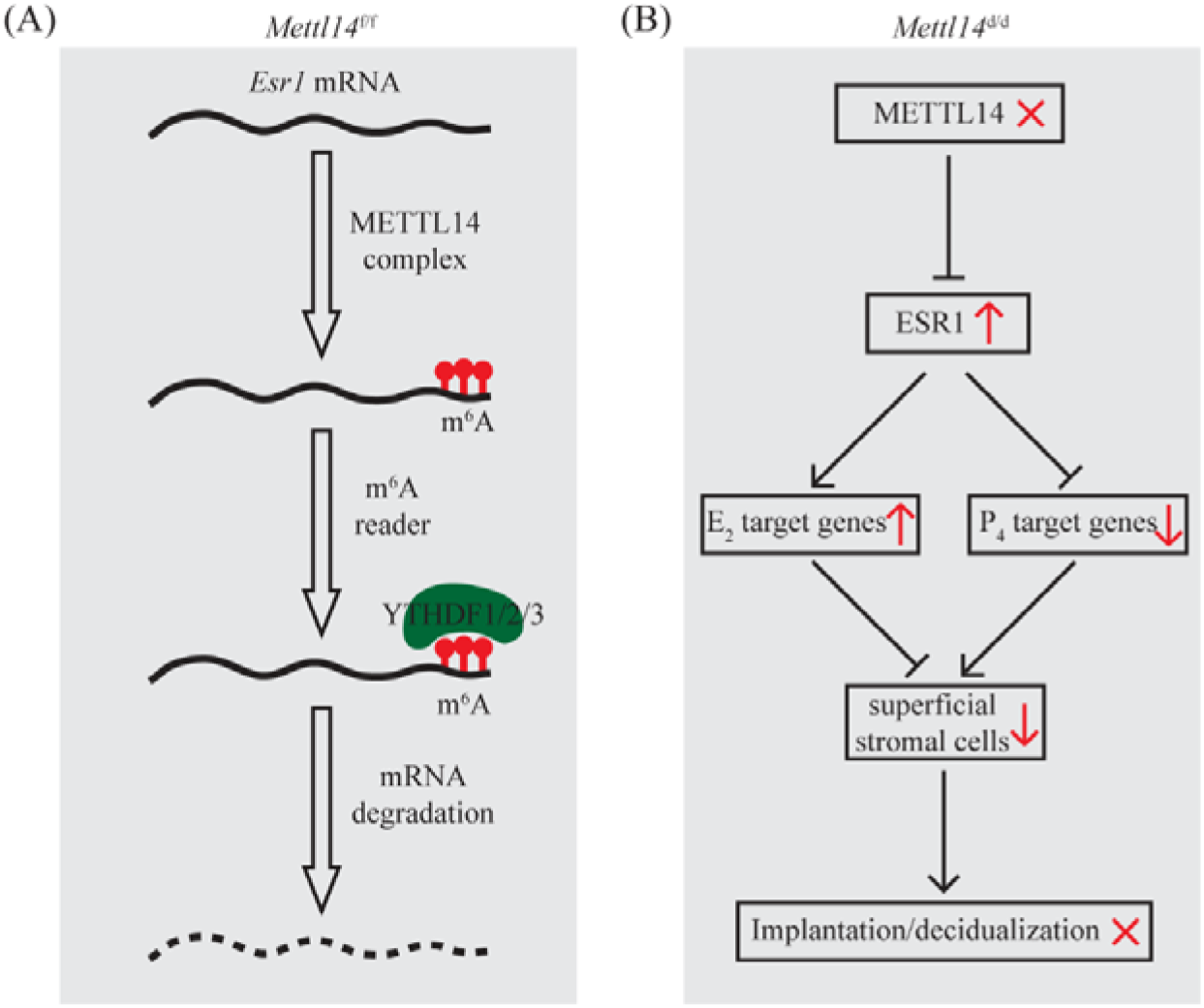
Working model. (A) In *Mettl14*^f/f^ mice, *Esr1* mRNA with m^6^A modification in the 3′-UTR is recognized by YTHDF1/2/3 to promote degradation. (B) In *Mettl14*^d/d^ mice, *Esr1* mRNA without m^6^A modification is resistant to degradation. High levels of ESR1 protein result in upregulation of E_2_ target genes, downregulation of P_4_ target genes and loss of superficial stromal cells, which eventually leads to failure in embryo implantation and decidualization.

The primary uterine phenotypes observed in *Mettl14*^d/d^ mice, namely complete failure of implantation and decidualization, are consistent with previous findings [22]. This phenotype is also analogous to that observed in uterine-specific *Mettl3* knockout mice [23, 24], given that METTL3 is another key component of the m^6^A writer complex. Using scRNA-seq, we found a substantial loss of SSCs in *Mettl14*^d/d^ mice. Notably, FGF1/2/7/10/22 are expressed in SSCs, while their corresponding receptor FGFR2 is expressed in luminal epithelial (LE) and glandular epithelial (GE) cells. The FGF pathway is crucial for establishing uterine receptivity in LE cells [4]. SSCs express high levels of FST, HOXA10, HOXA11, and HAND2 [12–15], all of which are essential for implantation and decidualization [4, 25–29]. Taken together, we propose that the loss of SSCs represents a key defect in *Mettl14*^d/d^ uteri, which may underlie the observed implantation and decidualization failure.

A previous study showed that *Mettl14* deletion caused evaluated E_2_ signaling via the ERK/AKT-pESR1 pathway [22]. In this study, we identified *Esr1* mRNA as a direct target of METTL14 in the uterus. We observed a marked increase in ESR1 expression in the uterus of *Mettl14*^d/d^ mice during the implantation window. Concurrently, E_2_ target genes such as *C3*, *Ltf*, *Muc1*, and *Muc4* were significantly upregulated. Notably, while PGR expression remained unchanged, P_4_ target genes like *Ihh*, *Areg*, *Hoxa10*, and *Hand2* were significantly downregulated. Elevated E_2_ activity in the uterus is known to suppress P_4_ target genes such as HAND2 [25–28], suggesting that increased E_2_ activity in *Mettl14*^d/d^ uteri may cause SSC loss.

As METTL3 and METTL14 form a heterodimer for RNA m^6^A methylation, it is usually assumed that their perturbations would yield similar defects. However, numerous studies have revealed that METTL3 and METTL14 have non-overlapping functions. For instance, *Mettl3* knockout showed embryonic lethality at E6.5 due to the inability to downregulate *Nanog* mRNA [30]. Conversely, *Mettl14* knockout mice exhibit abnormal embryo development from E6.5 onward, yet *Nanog* mRNA remains unaffected [20]. In cultured mouse embryonic stem cells, *Mettl3* knockout maintains a round and compact stem cell colony morphology, whereas *Mettl14* knockout was unable to sustain stemness [31]. In the mouse intestine, *Mettl14* deletion does not significantly impact the small intestinal epithelium [32, 33], but *Mettl3* deletion leads to epithelial distortion [34]. In mouse testis, the individual deletion of either *Mettl3* or *Mettl14* in advanced germ cells show normal spermatogenesis, but the combined deletion of both disrupts spermiogenesis [35]. These contrasting findings suggest that METTL3 and METTL14 function differently, potentially through their binding with a diverse array of partners. Previously, we and other researchers discovered that the conditional knockout of *Mettl3* in the mouse uterus resulted in embryo implantation failure due to a decrease in PGR expression at the protein level [23, 24]. Additionally, our studies in cultured mouse and human stromal cells confirmed the downregulation of PGR following *Mettl3* knockdown by siRNA [24]. Our mechanistic study revealed that METTL3-mediated m^6^A modification in the 5′-UTR of *Pgr* mRNA is imperative for its optimized translation efficiency in a YTHDF1-dependent manner [24]. However, in contrast to *Mettl3*^d/d^ uteri, in this study we observed that PGR expression remained unchanged in *Mettl14*^d/d^ uteri. Instead, there was an upregulation of ESR1 at the mRNA level. Our subsequent experiments using siRNA-mediated knockdown in cultured mouse and human stromal cells further validated these results. Mechanistically, we found that METTL14-mediated m^6^A modification in the 3′- UTR of *Esr1* mRNA enhances its degradation in a YTHDF1/2/3-dependent manner. Our findings, by highlighting the functional distinctions between METTL3 and METTL14 within the uterus, underscore the intricate and multifaceted role of m^6^A modification in governing uterine function.

In summary, by using the uterine-specific METTL14 deletion mouse model, we provide evidence that SSCs in the uterus are essential for embryo implantation.

## Materials and Methods

### Mice

*Mettl14*^f/f^ mice (Cat. No. NM-CKO-190007) and *Pgr*^Cre/+^ mice (Cat. No. NM-KI-200117) were purchased from Shanghai Model Organisms Center, Inc., China. *Mettl14*^d/d^ mice were produced by crossing *Mettl14*^f/f^ mice with *Pgr*^Cre/+^ mice. The *Mettl14*^f/f^ littermates were used as the wild-type control. Adult female mice were mated with normal fertile or vasectomized male mice to induce pregnancy or pseudo-pregnancy. The day of vaginal plug detection was designated as gestational day 1 (GD1). For embryo transfer experiments, 2-cell embryos were flushed from the oviduct of donor mice on GD2 and cultured in KSOM medium (Millipore) until they reached the fully expanded blastocyst stage. Six blastocysts were then transferred into one uterine horn of the pseudo-pregnant recipient mice on GD4, and uterine samples were collected on GD6. For artificial decidualization experiments, 20 μl of sesame oil (Sigma) was injected into one uterine horn of the pseudo-pregnant mice on GD4, and uterine samples were collected on GD8. All the animal procedures were approved by the Institutional Animal Care and Use Committee of South China Agricultural University (No. 2021B036).

### Primary culture of mouse uterine stromal cells

Mouse uterine stromal cells were isolated and cultured as previously described with minor modifications [36]. Briefly, uterine horns from wild-type mice on GD4 were dissected into small fragments. These tissues were then submerged in Hank’s balanced salt solution (HBSS) containing 6 mg/ml dispase and 25 mg/ml trypsin for 1 h at 4°C. Subsequently, the tissues were incubated for 1 h at room temperature and then for 10 min at 37°C. Following the removal of endometrial epithelial clumps, the remaining tissues were incubated once again in HBSS containing 0.5 mg/ml collagenase at 37°C for 30 min. To acquire stromal cells, the digested cells were filtered through a 70-μm mesh. The isolated cells were seeded onto 60- mm dishes at a density of 5 □ 10^5^ cells per dish and cultured in a mixture of phenol red-free Dulbecco′s modified Eagle′s medium and Ham F-12 nutrient medium (1:1; DMEM/F12; Gibco), supplemented with 10% charcoal-stripped fetal bovine serum (C-FBS; Biological Industries) and antibiotics. After an initial 2-h incubation, the medium was replaced with a phenol red-free culture medium (DMEM/F-12, 1:1) containing 1% charcoal-stripped FBS, 10 nM E_2_, and 1 μM P_4_ to induce in vitro decidualization.

### Quantitative RT-PCR

Total RNA was extracted using the TRIzol reagent (Invitrogen), and 1 μg of this purified RNA was used to synthesize cDNA with the HiScript □ RT Super Mix (Vazyme). Quantitative PCR was performed using the ChamQ SYBR qPCR Master Mix (Vazyme), executed on the Applied Biosystems 7300 Plus (Life Technologies). data were normalized against the expression of *Rpl7* as a reference gene. The complete list of primer sequences used in this study is provided in **Table S4**.

### Western blot

Uterine tissues and cultured cells were lysed using Radio-Immunoprecipitation Assay (RIPA) buffer supplemented with protease inhibitors (Roche). Total protein extracts were then resolved on a 10% SDS-PAGE gel and transferred onto a nitrocellulose membrane (Millipore).

Prior to antibody probing, the membrane was blocked with 5% skim milk in TBST.

Subsequently, the membrane was incubated overnight at 4°C with the primary antibody. Following thorough washing, the membrane was incubated with an HRP-conjugated secondary antibody for 1 h at room temperature. The immunoreactive signals were visualized using the ECL chemiluminescent kit (Amersham Biosciences). ACTB and GAPDH bands were used as loading control. Band intensities were quantitatively analyzed using the ImageJ software v1.54i [37]. Primary antibodies used in this study are listed in **Table S5**. Uncropped and unprocessed scans of the blots are presented in **Fig. S6**.

### Dot blot

The mRNA was harvested from total RNA using the Dynabeads^®^ mRNA Direct^TM^ Purification Kit (Invitrogen). After denaturing the RNA samples at 95 °C, they were immediately chilled on ice. Subsequently, the RNA samples were spotted onto an Amersham^TM^ Hybond^TM^-N+ membrane (GE Healthcare) and affixed to the membrane through UV cross-linking. To reduce non-specific binding, the membrane was blocked with 5% non-fat milk for 1 h. Following this, the membrane was incubated overnight at 4°C with an m6A-specific antibody. After thorough washing with PBST, the membrane was incubated with the secondary antibody, HRP-conjugated goat anti-rabbit IgG, for 1 h at room temperature. The immunoreactive signals were visualized using an ECL chemiluminescent kit (Amersham Biosciences). Methylene blue staining was performed as a loading control. Dot intensities were quantitatively analyzed using the ImageJ software v1.54i [37].

### Immunohistochemistry

Paraformaldehyde-fixed paraffin-embedded uterine tissues were sliced into 5-μm sections. Heat-induced antigen retrieval was then conducted in a 10 mM citrate buffer (pH = 6.0) for a duration of 15 min. To eliminate endogenous peroxidase activity, the sections were treated with 3% H_2_O_2_ for 20 min. After blocking with 10% horse serum in PBS, sections were incubated with the primary antibody overnight at 4°C. After this, they were incubated with an HRP-conjugated secondary antibody for one hour at room temperature. The immunoreactive signals were visualized using DAB staining kit (Zhongshan Golden Bridge Biotechnology Co., Beijing). For contrast, the sections were counterstained with hematoxylin. Primary antibodies used in this study are listed in **Table S5**.

### Alkaline phosphatase staining

Frozen tissue sections, precisely sliced at 10 μm, were swiftly fixed in cold acetone for 15 minutes and subsequently washed thoroughly three times with PBS. To visualize alkaline phosphatase activity, the BCIP/NBT kit (Zhongshan Golden Bridge Biotechnology Co., Beijing) was employed, following the manufacturer’s recommended protocol. For counterstaining, a 1% solution of methyl green was utilized.

### Scanning electron microscopy

To expose the luminal epithelium, the mouse uterus was carefully sliced open longitudinally. The samples were then gently washed with PBS and promptly fixed in 2.5% glutaraldehyde. Following this initial fixation, the samples underwent three washes in 0.1 M phosphate buffer (PB, pH = 7.4) and were subjected to a secondary fixation step using 1% osmium tetroxide for 1h at room temperature. After another three washes in 0.1 M PB (pH = 7.4), the samples were dehydrated through a gradually increasing series of ethanol concentrations and dried using a critical point dryer (Quorum, K850). The dried samples were then affixed to metallic stubs using carbon stickers and coated with a thin layer of gold for 30 s using an ion-beam sputtering apparatus (HITACHI, MC1000). Finally, the prepared samples were observed under a scanning electron microscope (HITACHI, SU8100) at an accelerating voltage of 3.0 kV.

### Transmission electron microscopy

The mouse uterus was initially fixed in 2.5% glutaraldehyde and then post-fixed with 1% osmium tetroxide. Following three thorough washes in 0.1 M PB (pH = 7.4), the samples underwent a graded dehydration process through various concentrations of ethanol, followed by infiltration with acetone. The infiltrated samples were then embedded in Embed 812 resin (SPI, 90529-77-4). Utilizing an ultra-microtome (Leica, UC7) equipped with a diamond slicer (Diatome, Ultra 45°), the resin blocks were precisely cut into 80-nm sections. These sections were subsequently stained with a 2% uranium acetate solution saturated in alcohol, followed by a 2.6% lead citrate solution. Finally, the stained samples were observed under a transmission electron microscope (HITACHI, HT7800) at an accelerating voltage of 80 kV.

### RNA-seq

Uterine segments were frozen in liquid nitrogen and stored at -80 °C until use. Total RNA extraction, cDNA library construction and high-throughput DNA sequencing were executed following our previously established protocols [38]. After quality control, the sequence data were aligned to the mouse genome (UCSC mm10) using HISAT v2.2.1 [39]. Gene-level expression values were derived from the aligned sequences utilizing Cufflinks v2.2.1 [40]. Differentially expressed genes were selected based on the following criteria: fold change > 2 and adjusted P-value < 0.05. To gain insights into the functional roles of differentially expressed genes, gene ontology analysis was performed using the DAVID online tools v2024q1 [41], with an adjusted P-value cutoff set at 0.05. Redundant enriched terms were manually eliminated.

### Single-cell RNA-seq

Uterine tissues were minced with a blade on ice. Single-cell library construction using the 10× Genomics protocol and high-throughput sequencing were performed according to our previous studies [12, 13, 15]. The gene expression matrix was computed using the CellRanger software (v3.0.1, 10× Genomic) and was further processed with the R package Seurat v3.1.3 [42]. For quality control, both cells with fewer than 200 or greater than 6000 unique genes and cells with greater than 25% of mitochondrial counts were excluded. Filtered data were normalized and subjected to principal component analysis (PCA) to reduce the dimensionality of the data. The JackStraw procedure was employed to determine the optimal number of PCA components. Utilizing the K-nearest neighbor (KNN) graph algorithm in PCA space, cells were clustered and visualized through a t-distributed stochastic neighbor embedding (tSNE) plot. The cell-type label for each cell cluster in the tSNE plot was manually assigned based on canonical markers. The CellChat v2.0.0 software [43] was used to infer cell-cell communication based on ligand-receptor interactions.

### MeRIP-seq

Purified mRNA was fragmented in fragmentation buffer (Invitrogen, AM8740) by heating it at 94□ for 5 min. The m^6^A antibody immunoprecipitated RNA fragments, along with the input control, were prepared according to our previous study [24]. MeRIP-Seq libraries were generated with the Next® Ultra™ II Directional RNA Library Prep Kit (NEB) and sequenced on the Illumina HiSeq 2500 system. After quality control, the sequence data were aligned to the mouse genome (UCSC mm10) with HISAT v2.2.1 [39]. For visual inspection, the sequence alignment on the genome was displayed using the IGV tool v2.14.0 [44]. Peak calling was performed with MACS software v3.0.0a7 [45] with q-value < 0.05. The metagene profile was analyzed using the R package Guitar v2.12.0 [46]. Motif analysis was carried out with HOMER v4.7 [47].

### Luciferase reporter assay

A segment of the mouse Esr1 3′-UTR (1-1539 bp) was synthesized and cloned into the psiCHECKTM-2 vector (Promega) by using the multiple cloning sites at the 3′-UTR of the Renilla luciferase gene. Three single-point mutants, A396T, A1005T and A1459T, as well as a triple-point mutant A396T/A1005T/A1459T, were generated using a QuickMutation™ Site- Directed Mutagenesis Kit (Beyotime Biotechnology, Shanghai). Reconstructed plasmids were transfected into HEK293T cells using Lipofectamine 3000 (Invitrogen). Cell lysates were collected 48 h after transfection. Luciferase activity in cell lysates collected 48 h after transfection was measured using the Dual-Luciferase Reporter Assay System (Promega). The siRNAs used in this study are listed in **Table S6**.

### RNA decay assay

Primary mouse and human endometrial stromal cells were plated in 12-well dishes and incubated overnight at 37 °C. For investigating METTL14-mediated mRNA decay, the cells were initially transfected with either a control siRNA or a specific *Mettl14* siRNA. Following transfection, the cells were treated with actinomycin D (A1410, Sigma) at a final concentration of 5 μg/ml to halt transcription. At various time intervals, total RNA was isolated from the cells. Subsequently, 50 ng of the isolated total RNA was subjected to quantitative RT-PCR to quantify the expression levels of *Esr1* mRNA. The obtained data were normalized to the initial time point (t = 0) for comparison. The siRNAs used in this study are listed in **Table S6**.

### Isolation and culture of primary HESCs

Three healthy fertile participants, who exhibited no signs of endometrial pathology and had a clinically confirmed pregnancy following embryo transfer, were selected for this study. Prior to commencing the study, it was approved by the Ethical Committee of the First Affiliated Hospital of Sun Yat-sen University (Approval No. 2018-266), and all participants provided their informed consent. The detailed characteristics of these patients are summarized in **Table S7**. The endometrial tissues were dissected into pieces and then underwent digestion using type I collagenase (Gibco) for 1 h. Subsequently, the endometrial epithelial cells and stromal cells were separated using membrane filters (100 µm and 40 µm cell filters, Corning). The human endometrial stromal cells (HESCs) were cultured in DMEM/F12 medium (Gibco), supplemented with 10% charcoal-stripped fetal bovine serum (cFBS, VivaCell). To induce decidualization, the cells were treated with 0.5 mM 8-Br-cAMP (Sigma) and 1 μM medroxyprogesterone acetate in 2% cFBS for 4 days.

### Statistical analysis

Statistical analysis was performed using R v4.1.3 (https://www.r-project.org/). The two-tailed unpaired Student’s t test was used for comparison of means of two groups. The one-way ANOVA with Tukey’s multiple comparison test was used for comparison of means of more than two groups. Data are presented as the mean ± SD and P < 0.05 was considered statistically significant.

## Acknowledgements

This research was funded by National Natural Science Foundation of China (32370913 and 32400710), Guangdong Natural Science Funds for Distinguished Young Scholars (2021B1515020079), Innovation Team Project of Guangdong University (2019KCXTD001), Guangdong Special Support Program (2019BT02Y276), National Key R&D Program of China (2018YFA0801404), and Double first-class discipline promotion project (2023B10564003).

## Author contributions

Y.-W.X., S.-H.Y. and J.-L.L. designed research; Z.-H.Z., R.-F.J., Y.-Q.H., Q.-Y.Z., Y.-T.D., J.- Q.Z. and J.-P.H. performed the experiments; Z.-H.Z., C.-H.D., Y.-W.X., S.-H.Y. and J.-L.L. analyzed the data. Z.-H.Z., Y.-W.X., S.-H.Y. and J.-L.L. wrote the paper. All authors read and approved the final paper.

## Data availability statements

The sequencing data generated in this study are deposited in the Gene Expression Omnibus (GEO) under accession codes GSE263378, GSE263379 and GSE264490.

