## Supplementary figures and tables for "Superficial stromal cell population in the mouse uterus require METTL14 for development and functional competence to support embryo implantation"

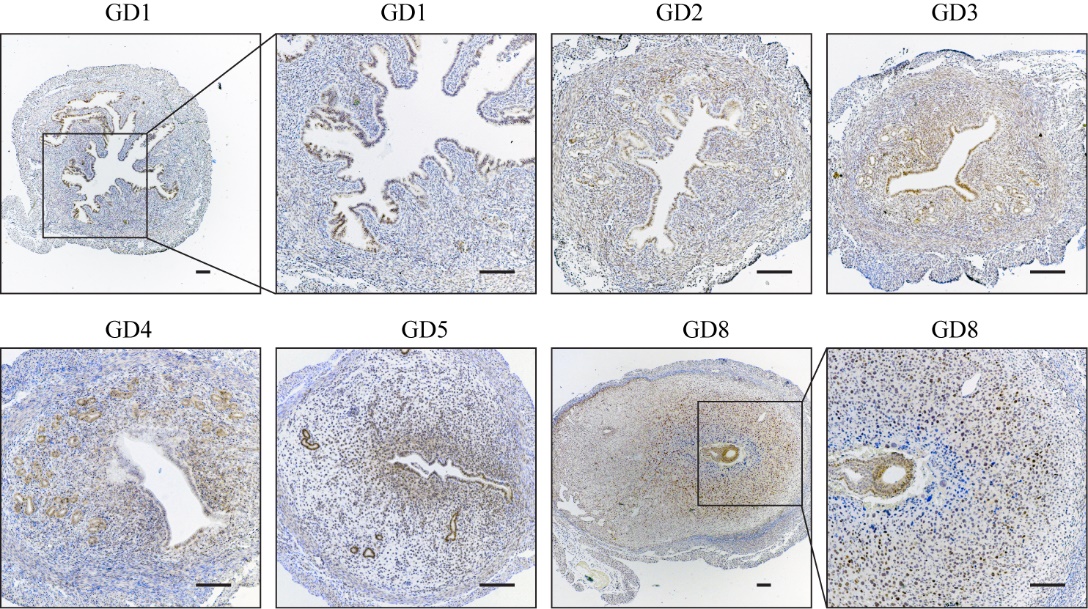

**Supplementary Figure 1. Immunohistochemical staining of METTL14 protein in wild-type mouse uterus during the peri-implantation period.** GD, gestational day. GD1 is the day when the vaginal plug is seen. Scale bar = 100 μm.

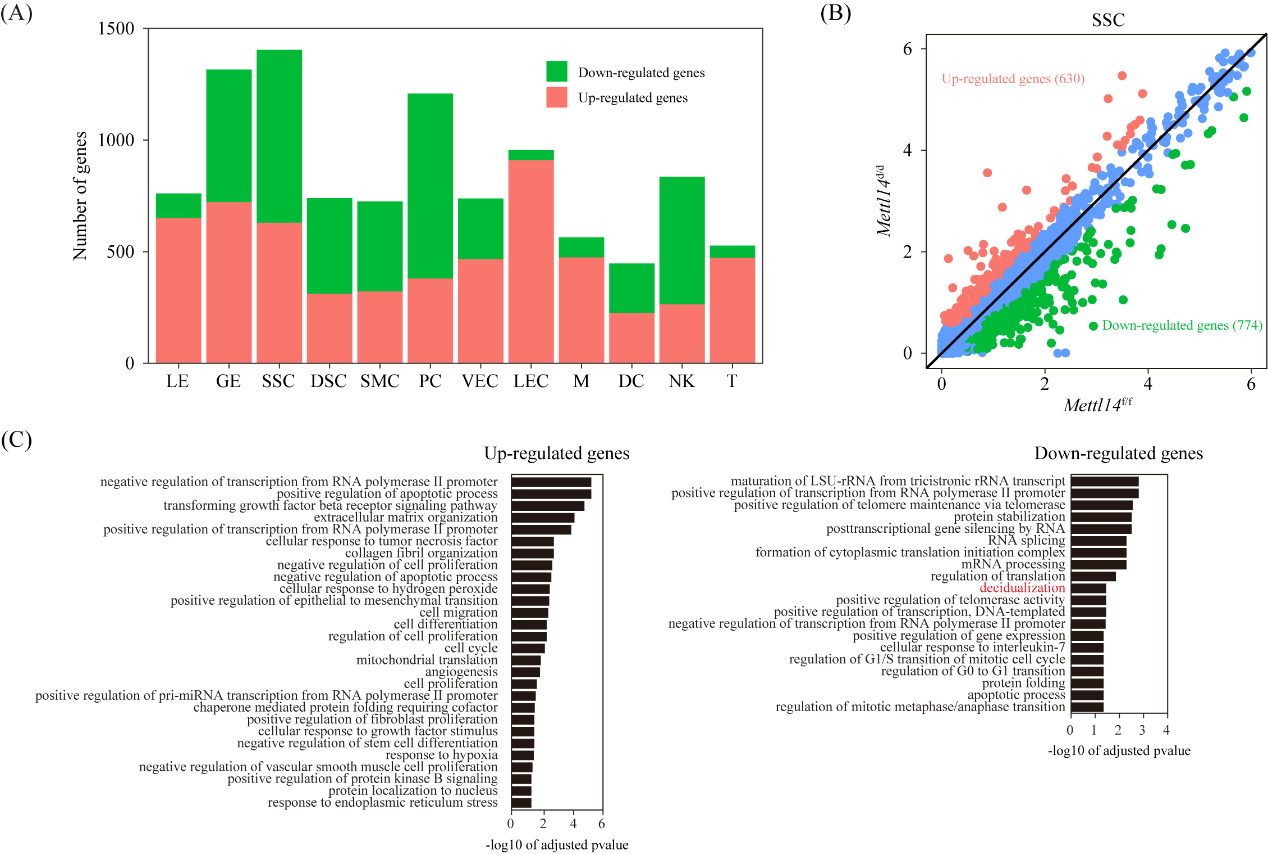

**Supplementary Figure 2. Differentially expressed genes in each cell type in *Mettl14*^d/d^ uteri compared *Mettl14*^f/f^ uteri based on single-cell RNA-seq.** (A) Bar plot showing the count of differentially expressed genes in each cell type. The threshold values for differentially expressed genes were fold change > 1.5 and P < 0.05. LE, luminal epithelial cells; GE, glandular epithelial cells; SSC, superficial stromal cells; DSC, deep stromal cells; SMC, smooth muscle cells; PC, pericytes; VEC, vascular endothelial cells; LEC, lymphatic endothelial cells; NK, natural killer cells; T, T cells; M, macrophages; DC, dendritic cells. (B) Scatter plot for the comparison of gene expression levels in SSCs between *Mettl14^d/d^* uteri and *Mettl14^f/f^* uteri. Non-changed genes were shown in blue color, while differently expressed genes were denoted in red or green. (C) Gene ontology enrichment analysis of differentially expressed genes in SSCs conducted by using the DAVID online tools. The significance threshold for adjusted P value was set at 0.05.

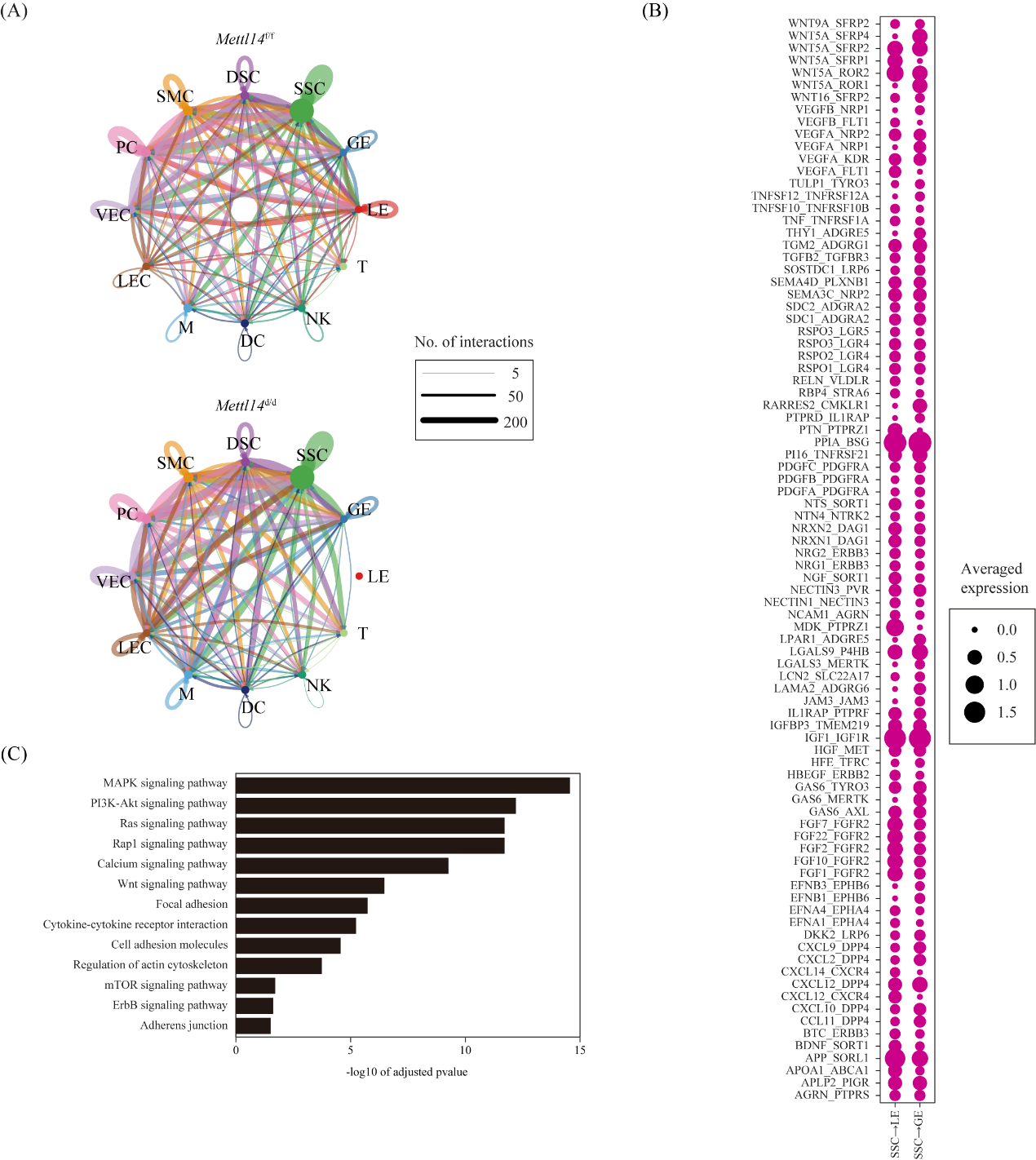

**Supplementary Figure 3. Cell-cell communication between different cell types based on single-cell RNA-seq data.** (A) Network plot showing ligand-receptor interactions underlying the cross-talk between different cell types. The number of interactions was indicated by the degree of thickness. (B) Dot plot showing selected ligand-receptor interactions underlying the cross-talk between SSCs and LE/GE. Only interactions with an averaged expression level greater than 0.1 was shown. (C) KEGG Pathway enrichment analysis of ligand-receptor pairs performed by using the DAVID online tools. The significance threshold for adjusted P value was set at 0.05.

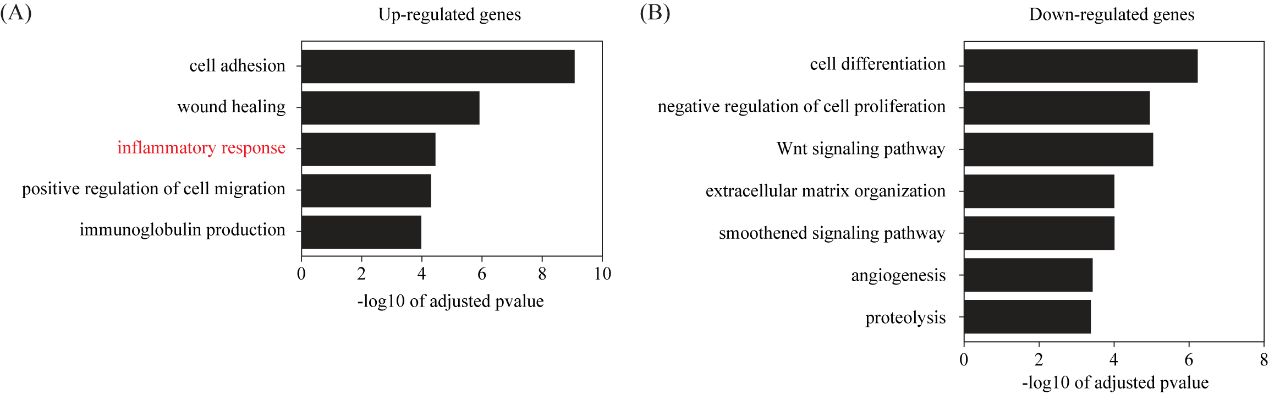

**Supplementary Figure 4. Gene ontology enrichment analysis of up-regulated genes (A) and down-regulated genes (B) in the *Mettl14^d/d^* uteri compared with the *Mettl14^f/f^* uteri on GD4 based on RNA-seq.**

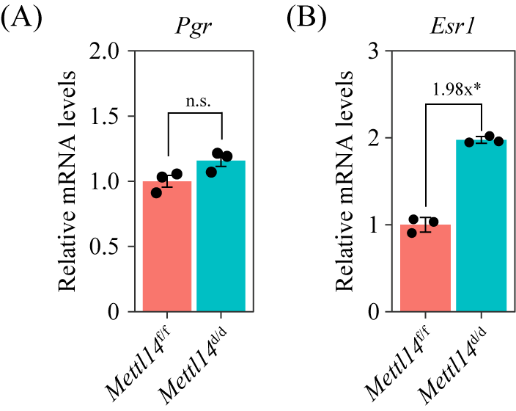

**Supplementary Figure 5. Quantitative RT-PCR analysis of *Pgr* (A) *and Esr1* (B) mRNA levels in the uterus of *Mettl14*^f/f^ and *Mettl14*^d/d^ mice on GD4.** Data are presented as mean ± SD. *, P < 0.05.

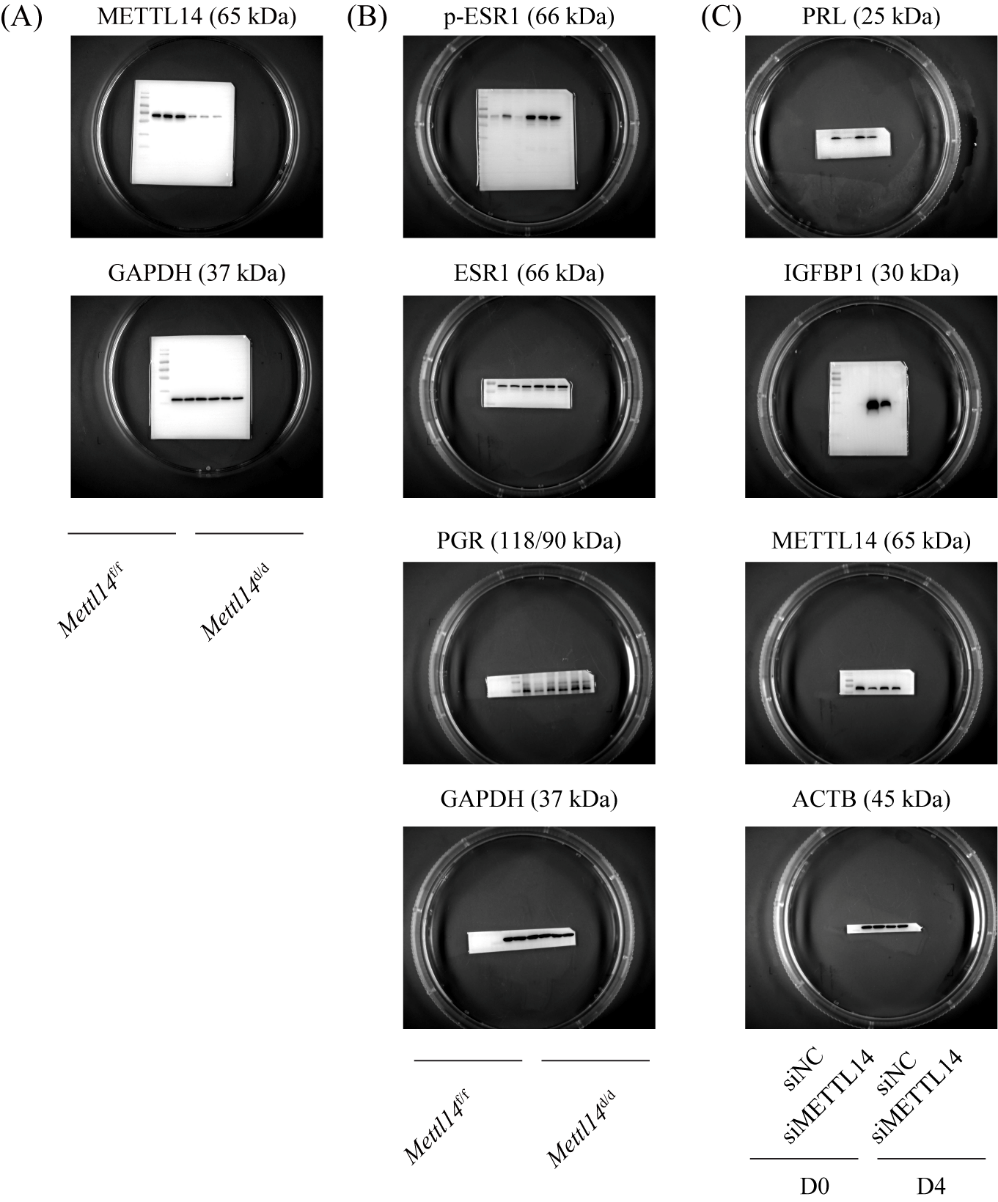

**Supplementary Figure 6. Pictures of the uncropped Western blots.** (A) Fig. 1E. (B) Fig. 5C. (C) Fig. 8D.

**Supplementary Tables**

**Supplementary Table 1. Differentially expressed genes in *Mettl14*^d/d^ compared to *Mettl14*^f/f^ uteri on GD4 identified by single-cell RNA-seq (fold change > 1.5 and adjusted p-value < 0.05).**

**Supplementary Table 2. Differentially expressed genes in *Mettl14*^d/d^ compared to *Mettl14*^f/f^ uteri on GD4 identified by bulk-tissue RNA-seq (fold change > 2 and adjusted p-value < 0.05).**

**Supplementary Table 3. The complete list of m^6^A peaks in wild-type mouse uterus on GD4 identified by MeRIP-seq (q-value < 0.05).**

**Supplementary Table 4. Primers used in this study.**

**Supplementary Table 5. Antibodies used in this study.**

**Supplementary Table 6. siRNAs used in this study.**

**Supplementary Table 7. Detailed information of human participants for isolation of primary endometrial stromal cells in this study.**
